# Cranes soar on thermal updrafts behind cold fronts as they migrate across the sea

**DOI:** 10.1101/2023.06.29.547013

**Authors:** Sasha Pekarsky, David Shohami, Nir Horvitz, Rauri C. K. Bowie, Pauline L. Kamath, Yuri Markin, Wayne M. Getz, Ran Nathan

## Abstract

Thermal soaring conditions above the sea have long been assumed absent or too weak for terrestrial migrating birds, forcing large obligate soarers to take long detours and avoid sea crossing, and facultative soarers to cross exclusively by costly flapping flight. Thus, while atmospheric convection does develop at sea and is utilized by some seabirds, it has been largely ignored in avian migration research. Here we provide direct evidence for routine thermal soaring over open sea in the common crane, the heaviest facultative soarer known among terrestrial migrating birds. Using high-resolution biologging from 44 cranes tracked across their transcontinental migration over 4 years, we show that soaring characteristics and performance were no different over sea than over land in mid-latitudes. Sea-soaring occurred predominantly in autumn when large water-air temperature difference followed mid-latitude cyclones. Our findings challenge a fundamental paradigm in avian migration research and suggest that large soaring migrants avoid sea crossing not due to absence or weakness of thermals but due to their uncertainty and the costs of prolonged flapping. Marine cold air outbreaks, imperative to the global energy budget and climate system, may also be important for bird migration, calling for more multidisciplinary research across biological and atmospheric sciences.

## Introduction

Sea crossing is highly challenging for migrating terrestrial soaring birds that regularly soar and glide over land (1, 2), leading to interspecific variation in sea-crossing strategies explained by wing morphology and flight modes (3; Fig. 1). Obligate soaring birds take long detours to circumvent the sea (4-6) or cross it only in short sections (7). The prevailing paradigm in avian migration research assumes that such detours are caused by the absence or weakness of thermal soaring conditions above the sea. Most terrestrial soaring birds are facultative soarers that are also capable of prolonged flapping and therefore, under the same paradigm, have been assumed to cross the sea exclusively by costly flapping flight (1).

**Figure 1.**
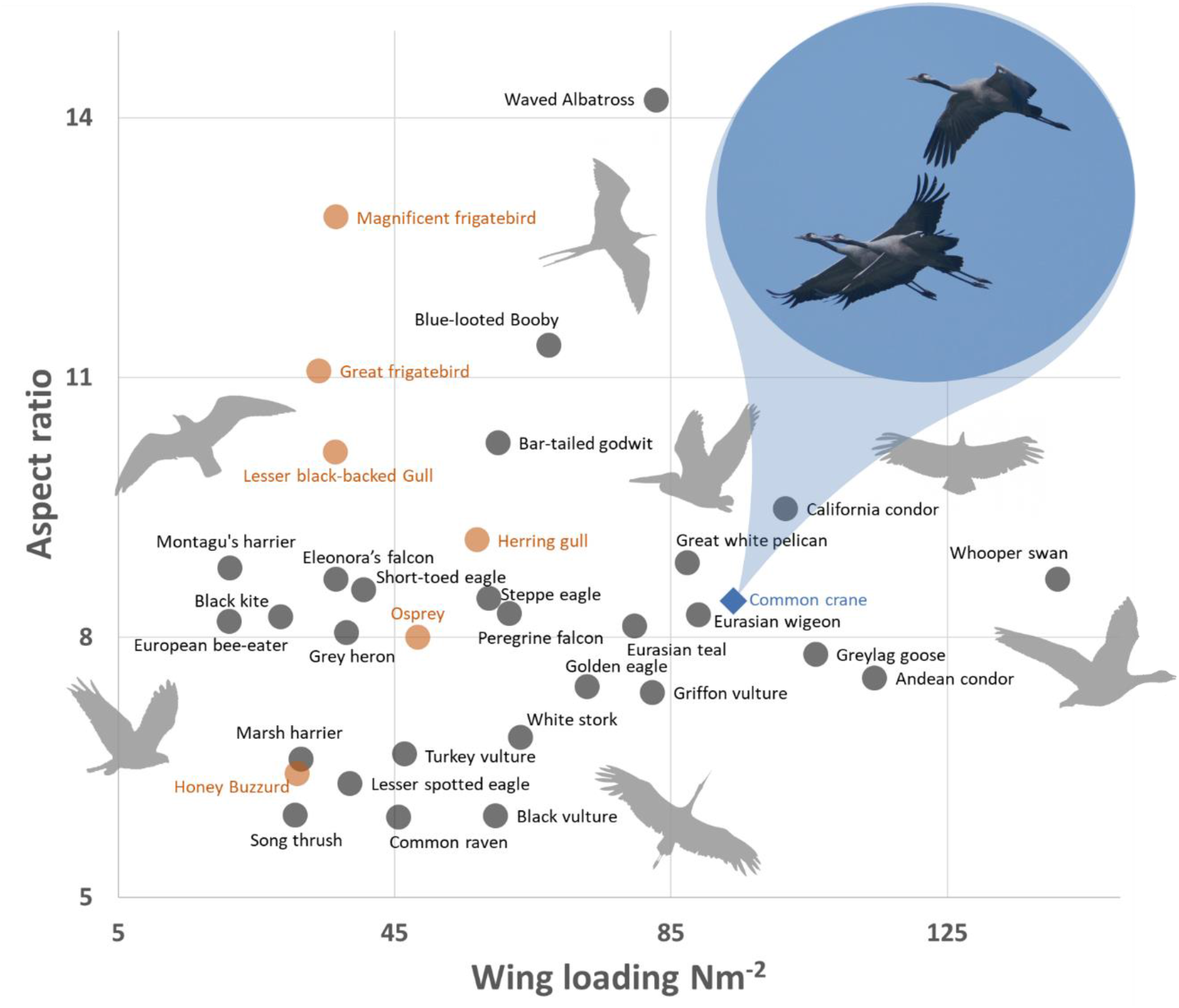
Morphospace defined by the relationship between wing loading and aspect ratio. The common cranes (blue) positioned in the morphospace near obligatory soarers such as vultures and obligatory flappers such as geese (morphological data in SI Appendix, Table S1). Species shown or suspected to perform thermal soaring at sea (orange) generally have lower wing loading compared to cranes. Photo: R.N.

Nonetheless, ascending air through atmospheric convection does develop over oceans and seas, encompassing atmospheric phenomena such as radiative cooling above cloud tops, trade-wind convection over tropical waters, and marine cold air outbreaks outside the tropics (8-10). In the latter, convection is generated when colder continental air flows over warmer sea-surface water. Under such temperature difference (ΔT; positive if sea surface is warmer than the overlying air), the cold air layer is heated by the water, producing unstable upward-downward air motion in the marine atmospheric boundary layer that may organize into convective cells (8, 9). Recent studies have shown how the unique combination of trade-wind convection over tropical oceans and extreme wing morphology of frigatebirds enables them to soar as they forage for extended periods (11), and that gulls use soaring flight when foraging at sea relatively close to shore(12). Yet, despite very early observations of soaring gulls that inspired the meteorological and physical study of cellular convective systems over the oceans (13, 14), their utilization for soaring by terrestrial birds over temperate and subtropical waters has been largely ignored in avian migration research (1, 15, 16) and the paradigm uncontested.

Recent studies have either suggested thermal soaring during sea crossing based on low-resolution tracking of raptors (17) or examined uplift potential during raptor migration over the sea (18-20), but lacked direct biologging evidence for the circling behavior typical of thermal soaring. Consequently, sea crossing solely by flapping flight could not be excluded in these studies. The first, and so far only, direct evidence based on high-resolution GPS tracking comes from ospreys migrating across the Mediterranean Sea (21), which have a relatively low morphospace position (Fig. 1). Moreover, the underlying meteorological conditions responsible for generating uplift potential that can be (and are) used by migrating birds remain largely under-investigated both in ecological and meteorological studies.

Among migratory birds, cranes are the heaviest facultative soarers that commonly use both low-cost soaring and costly prolonged flapping flight when and where soaring is unfeasible (1, 22). Cranes have a unique combination of wing morphology traits that neither fit terrestrial nor marine facultative soarers, but lies well within a zone merging the largest obligate soarers (pelicans, vultures and condors) that seldom or never cross the sea, and the largest obligate flapping birds (swans and geese) that use prolonged flapping for sea crossing (Fig. 1). Common cranes (*Grus grus*) routinely cross the Baltic, Black, Mediterranean and Red Seas during migration, including long stretches of up to 850 km over open water (22, 23). The common and the white-naped cranes (*Antigone vipio*) are the heaviest facultative soarers known to migrate long distances across the sea and, as in other cases, have been assumed to do so by combining gliding and flapping (1, 24), but not by thermal soaring.

Here, we used 1524 hours of high-resolution (1Hz) GPS, three-dimensional acceleration and magnetometer measurements from 44 common cranes, tracked along the breadth of their cross-continental migratory route between western Russia and Africa during 2018-2021, to provide the first direct evidence that cranes repeatedly use thermal soaring over the sea and far from nearest land (Fig. 2). Since cranes are terrestrial flight generalists and are thus expected to better exploit the environmental heterogeneity available at their aerial habitat (3, 25), we analyzed soaring and flapping flight performance and characteristics to investigate their sea crossing mechanisms, as well as the underlying large scale meteorological conditions that enable them to soar and glide over extended seascapes.

**Figure 2.**
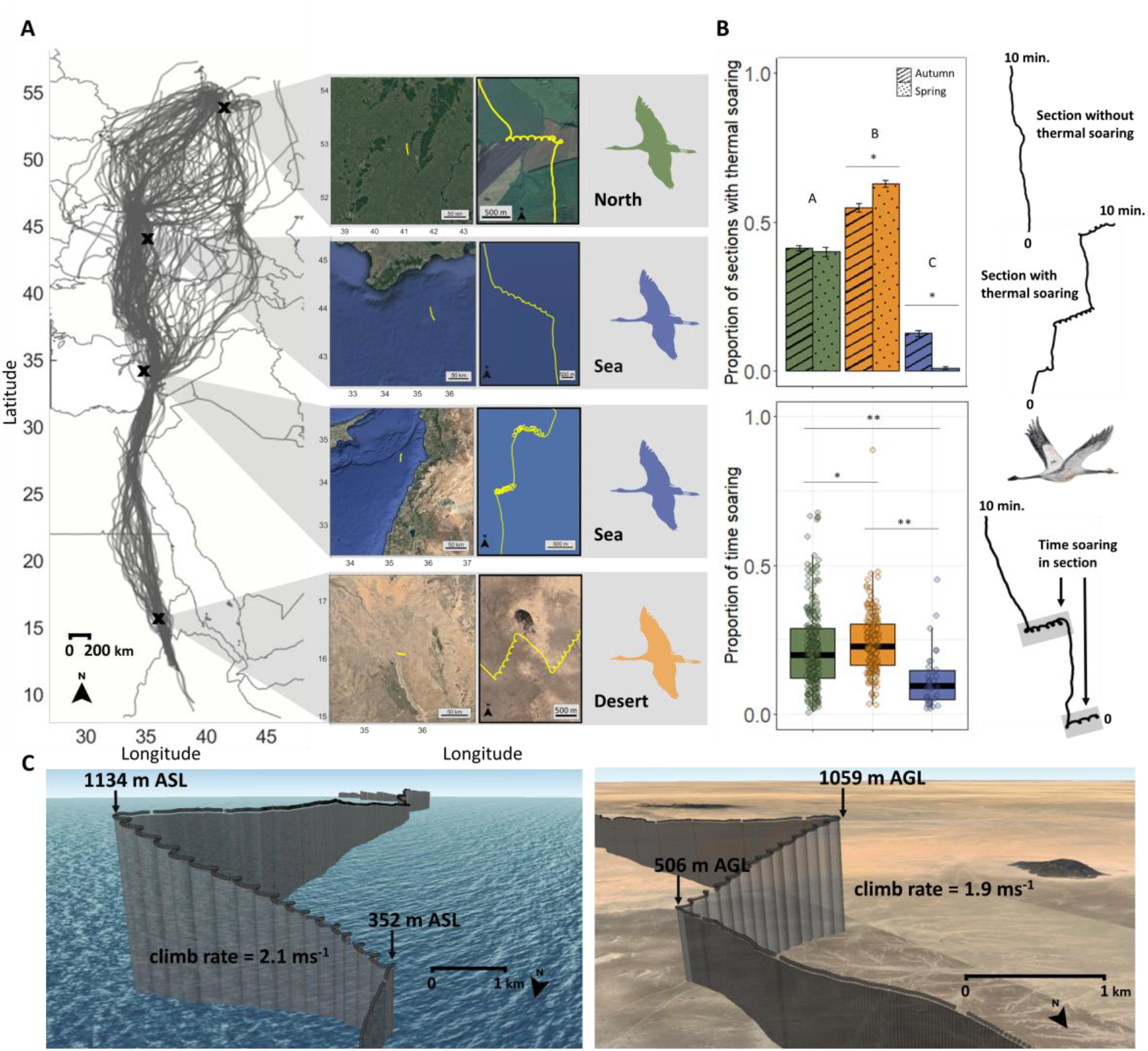
Soaring during migration trips over sea and land. (A) GPS tracks of 44 cranes for 135 autumn and spring migrations. To the right, examples of high temporal resolution (1-Hz) 10-min sections with soaring over land >32N (green, “north”), the Black and Mediterranean seas (blue), and land <32N (orange, “desert”). (B) Top: proportion of 10-min. sections with thermal soaring at different geographical regions. Asterisks and letters indicate significant difference between seasons and geographical regions, respectively (χ2, Marascuilo post-hoc, α=0.05; Table S2, S3); Bottom: proportion of time soaring within 10-min. sections during migration days with at least one soaring event. Asterisks indicate significant difference between regions (* p<0.05, ** p<0.001; ART-ANOVA with pairwise Bonferroni post-hoc). (C) Three-dimensional view of a migratory track segment in 1-Hz above the Black Sea (left) and above the Sahara Desert in Sudan (right).

## Results

High-resolution biologging of tracked cranes was configured in designated areas over three regions: land north of 32N, sea (Black and Mediterranean), and land south of 32N (desert) (Fig. 2A, SI Appendix, Fig. S1). Consequently, the high-resolution dataset was divided into 8657 10-min sections, with 3412 sections containing at least one thermal soaring event. In at least 40% of the time, cranes migrated exclusively using flapping flight, even over land at southern latitudes (Fig 2B). The proportion of thermal soaring differed between regions (χ^2^=1373.2, d.f.=2, p<0.001) and seasons (χ^2^=1425.1, d.f.=5, p<0.001), with the highest over the desert and lowest over the sea (Fig. 2B, SI Appendix, Table S2), and lower during autumn than spring in the desert but much higher over the sea (Fig. 2B, SI Appendix, Table S3). In diurnal trips with soaring activity, the proportion of time soaring was 2.3 times lower over sea than land (Fig 3B). The mean (±SD) air speed was 47.1±16.1 and 38.4±14.2 in sections with and without soaring, respectively (ART-ANOVA: F=201, p<0.001), indicating that soaring flight is likely to be favored not only for energy-but also for time-minimization, corroborating findings from large obligate soarers (27). Soaring-gliding performance over the sea was not different than over land at northern latitudes, but both were significantly different than over land at southern latitudes. Cranes tended to engage in more risk-prone flight (28) and have higher climbing rates, higher thermal exit height and lower flapping ratios in both the climbing and gliding phases, over southern latitude land (p<0.001; Fig 3, SI Appendix, Fig S3).

**Figure 3.**
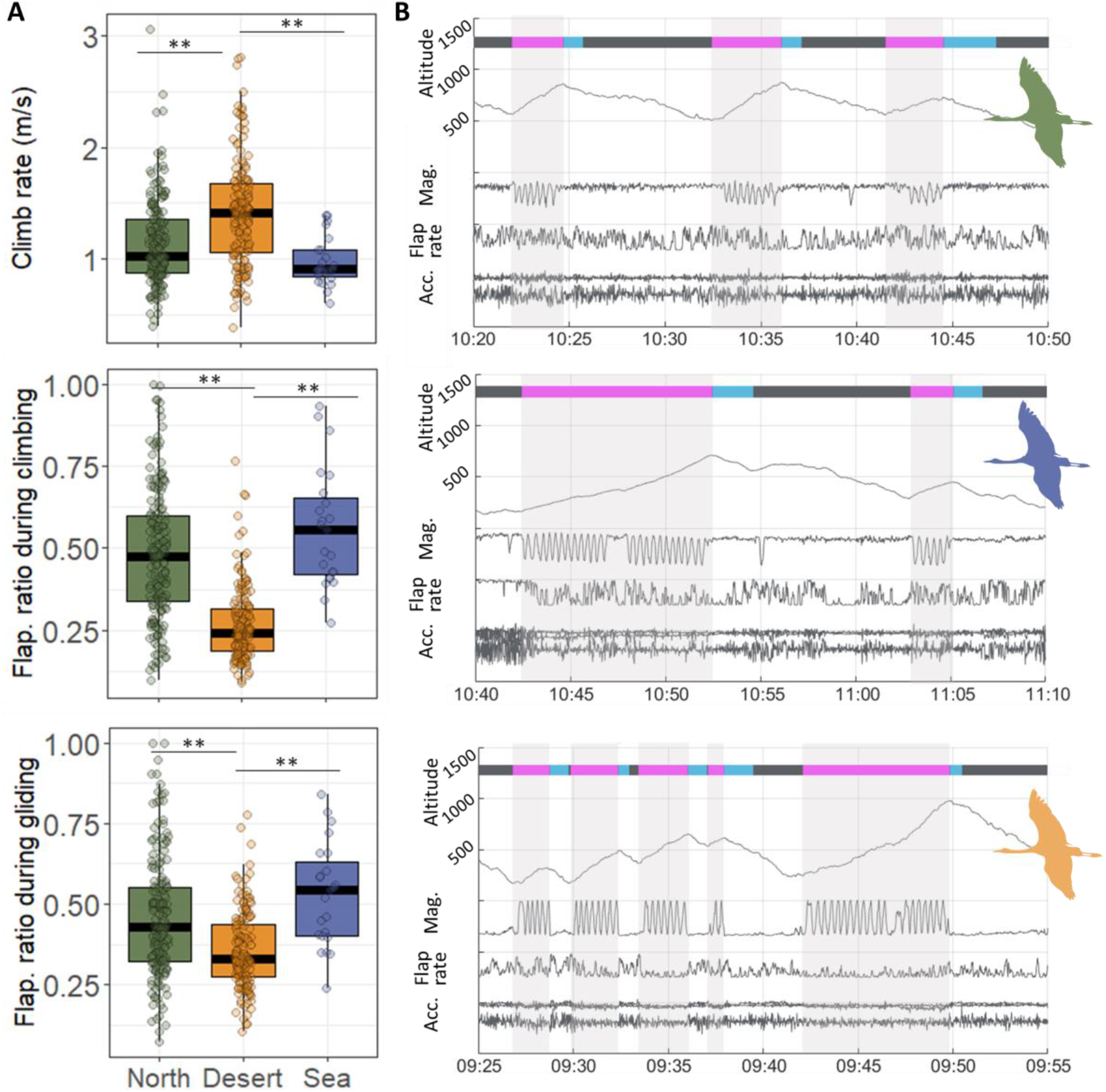
Soaring flight characteristics across three geographical regions. See Fig. 2 for region classifications and color codes. (A) Top: climb rate during soaring phase. Middle and bottom: flapping ratio during soaring and gliding phases, respectively, based on 1-Hz tri-axial acceleration. Boxplots represent averaged individual data of daily means (for days with more than one soaring-gliding events) with population median of all samples (north, N=182; desert, N=154; sea, N=23). Asterisks indicate significant difference between groups (* p<0.05, ** p<0.001; ART-ANOVA with pairwise Bonferroni post-hoc). (B) Typical ethograms of flight types within a single flight segment classified to circular thermal climbing (pink bar with grey shading) and gliding (blue bar) according to flight height (GPS) and heading change (GPS and magnetometer). Height above ground (m), magnetometer (x-axis; gauss), mean flap rate (flap s−1) and tri-axial acceleration shown (ms-2) over the north (top), sea (middle) and desert (bottom) regions. Time shown is UTC.

The probability of thermal soaring when crossing the sea was significantly and positively related to ΔT and wind speed at median flight height (395m), but wind speed effect was weak (Table SI Appendix, S4). Generally, thermal soaring occurred mostly in ΔT>1. To understand the broader meteorological context, we examined meteorological conditions 3 days before and 1 day after Mediterranean Sea crossing events during autumn (Fig. 4). For events including thermal soaring, all environmental conditions two and three days prior to departure were significantly different than the departure day (p<0.05, Fig. 4A blue lines). In contrast, sea crossing events that did not include soaring did not show clear trends in any of the meteorological variables (Fig. 4A red lines). ΔT levels were, on average, lowest 3 days before departure of soaring cranes (indicating relatively high air temperature very close to the sea temperature, p=0.02) and reached a maximum one day before departure. In contrast, crossings without soaring occurred during low ΔT levels without a sharp increasing trend in the preceding days (Fig. 4B). Additionally, crossings that included thermal soaring were preceded by a decrease, followed by a sharp increase, in sea level pressure, on average reaching a minimum two days before departure (p<0.05). They also occurred during a minimum in total cloud cover and daily precipitation, both reaching a maximum 2-3 days before departure (p=0.008 and p<0.001 respectively). Tailwind was present but weak during departure days of soaring cranes, but was strong headwind two and three days prior (p<0.05). Since autumn departures for Mediterranean Sea crossings are in a southerly direction (mean ± SD 285° ± 16), tail and headwinds generally correspond to northerly and southerly winds, respectively. The synoptic interpretation of these results indicates the passing of a mid-latitude cyclone (low-pressure area) and associated cold front two days, on average, prior to departures for Mediterranean Sea crossings that include thermal soaring (Fig. 4C).

**Figure 4.**
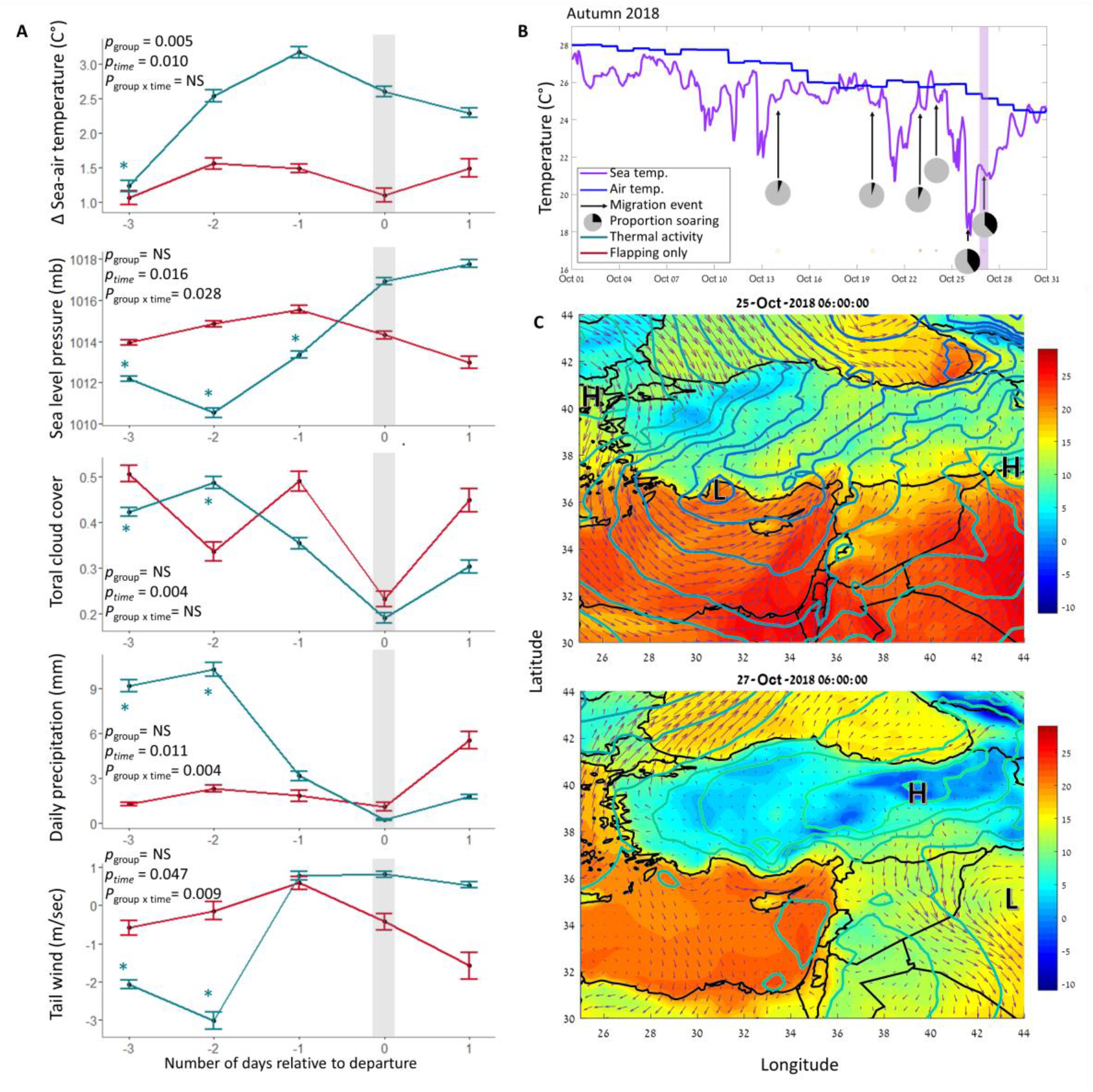
Synoptic analysis of Mediterranean Sea crossing events. (A) Time series of 4 days, with 0 being the day of departure from the stopover site in Turkey. Positive Δ sea-air temperature indicates cold air over a warm sea which is necessary for thermals to develop. Total cloud cover ranges between 0 (clear skies) and 1 (complete cloud cover). Tailwind is relative to travel direction. RM-ANOVA results shown of each metrological factor on bird flight mode (group) and timing (time) is shown. Asterisks denote significant (P<0.05) Dunnetts’ pairwise comparisons between a certain time and departure time. (B) Time-series of sea surface temperature (°C) in blue, and 2-m air temperature (°C) in purple exemplifying conditions during October 2018. Arrows indicate Mediterranean Sea crossing events; the corresponding pie charts show, in black, the percentage of crossing time that contained thermal soaring. Purple shading indicates the event for which weather conditions are shown in panel C. (C) Example of a low-pressure weather system affecting the eastern Mediterranean on October 25-27, 2018. Shaded areas are air temperature (°C) at 2-m height; colored lines are isobars of sea-level pressure (mb), with colder (bluer) colors indicating lower pressure; and arrows are wind vectors at an altitude of 950 mb (approximately 450 m a.s.l). “L” and “H” indicate low- and high-pressure areas, respectively. Top: October 25, 2018 (−2 timescales relative to departure), 0600 UTC: a relatively deep low-pressure area approaches the eastern Mediterranean from the west. Bottom: October 27, 2018 (departure day), 0600 UTC, the low-pressure area moved eastward and was replaced by high pressure over Turkey and a shallow insignificant low south-east of Cyprus.

## Discussion

Our data directly demonstrates thermal soaring over the sea by a large, heavy terrestrial migrant, with wing loading up to twice that of all raptors and gulls for which this behavior was previously documented (13, 14, 21) or suggested (17, 19, 20). Interestingly, we found that soaring crane climb rates over the sea were comparable to over land in northern latitudes, but the time spent soaring was considerably lower, suggesting lower frequency but not lower strength of thermals over the sea (Fig. 3). These findings challenge fundamental assumptions in avian migration research that have assigned a small, if any, role for thermals over the sea for migrating birds and explained sea avoidance of obligate soarers by thermal absence or weakness (2, 21, 29).

Our study shows the ability of facultative soaring migrants to switch flight modes in response to changes in the environmental conditions they encounter *en route*. Cranes heavily rely on powered flapping flight, use it exclusively for more than half of their migratory flights, and frequently flap also during thermal soaring-gliding (Fig. 3). Yet, unlike previous assertions that cranes use flapping as their primary flight mode and switch to thermal soaring when available to reduce flight energetic costs (22, 24), we found that soaring flight is also faster than flapping flight, hence likely favored also to minimize migration time, as shown in large obligate soarers (27). In addition, cranes merge soaring and flapping flight to “prolong” inter-thermal gliding, and to keep in tight flock formation also during thermal circling (1, 22). The higher climb rates and lower flap rates observed during migration over land at southern latitudes probably reflect stronger thermal activity in these areas while having few, if any, opportunities to refuel (1, 23).

Mediterranean sea-crossing events in autumn that included thermal soaring occurred on average two days after the passage of mid-latitude cyclones, when positive ΔT values occurred but the precipitation and headwinds associated with the cold front have ceased (Fig. 4). The low-pressure area draws in cold air behind it, which increases sea-air ΔT as the sea surface temperature is hardly affected by synoptic variations (Fig. 4B). Winds also tend to have a strong southerly component ahead of the low, which generates headwinds for the south-bound cranes. After the passing of the cold front and during gradual clearing of the low, the northerly (tail) component of the wind sets in, though cloudiness and precipitation may still linger. After that, even as ΔT decreases from its maximum, meteorological conditions are favorable for departure and for thermal soaring over the sea. During the spring, however, the same post-cyclonic cold air events generally correspond to headwinds for the north-bound cranes. This may partly explain why soaring over the Mediterranean was much rarer in spring.

In summary, we provide direct evidence based on rich high-resolution data for routine use of thermal soaring over the sea in the heaviest facultative soarer among terrestrial migrating birds, and investigate the mechanisms underlying this surprising finding, which calls to reconsider a prevailing paradigm in bird migration research. These mechanisms encompass the ability of cranes to adapt their flight mode to conditions *en route* and to flexibly select their migration route and timing, and the atmospheric convective processes that enable thermal soaring of such heavy birds over the sea. These atmospheric processes are imperative to understanding low cloud formation and the global energy budget and climate system (8, 9, 30). Here, we show that even small-scale marine cold air outbreaks such as those occurring in the eastern Mediterranean and Black Seas, which may not even generate visible convective cells (organized clouds) that are of interest to the atmospheric science community, still have significant global effects for biological processes such as bird migration and are of interest to the biological science community. Migratory birds can serve as sentinels of climatological and meteorological phenomena, sparking new opportunities for multidisciplinary research across biology and atmospheric sciences.

## Materials and Methods

### Tagging and data collection

Between January 2016 and September 2018, 44 common cranes (SI Data S1) were trapped in their pre-migration flocking areas in western Russia (Ryazan area; 54°56’N, 41°02E). The cranes were trapped using alpha-chlorolose (cf. 31, 32, 33) and processed in accordance with protocols approved by the Department of Environment of the Ryazan district, Russia (permit СК19-7154). Captured birds were color-ringed, fitted with leg-mounted solar-powered GPS-GSM transmitters (58 OrniTrack-L40: Ornitela, Lithuania), morphological measurements were taken, and body feathers were collected for molecular sexing. The maximal total mass of a transmitter plus rings used for attachment was (mean±SD) 0.8±0.09% (range: 0.7-1%; 35g-42g) of the captured cranes’ average body mass.

GPS locations were sampled at a resolution of 2 min to 1 h over the whole annual cycle depending on the measurement scheme and battery recharge (hereafter low-resolution dataset). Once a bird was flying inside pre-set geographical areas along the migratory route, GPS data was sampled continuously at 1 s intervals (hereafter high-resolution dataset; SI Appendix, Fig. S1). During 1Hz GPS recording, three-dimensional acceleration and magnetometer data were recorded in synchrony with the GPS position (Fig. 3B). Higher resolution (10Hz) 4-s bursts of three-dimensional acceleration and magnetometer were recorded once every 1 minute; during this ultra-high resolution sensor burst, GPS recording is paused. All data were downloaded remotely through a GSM network connection.

In all our analyses we used data from individuals migrating mainly along the Russian-Pontic route of the East Eurasian Flyway, leading from breeding grounds in Eastern Europe and the European part of Russia, through the Black Sea and towards wintering grounds in the Near East and north-east Africa (34). Smaller portion of individuals breeding in the European part of Russia also use the Caucasus flyway, leading across the Caucasus mountains to wintering grounds in Iran, the Near East and north-east Africa (23, 34). Age categories were assigned separately for every migration year, leading to some individuals being included in juvenile category during the first year of data collection and later assigned to adult category. Breeding adults were defined as adult birds observed with juveniles in current or previous years. All movement data analyses were performed using MATLAB R2020a (The Mathworks Inc., Natick, MA, USA).

### Movement analysis

High-resolution (1Hz) data was recorded for various lengths of time depending on battery recharge and point of entry/exit to/from the predefined preset geographical areas. We included in the analysis only continuous 1Hz sections longer than 10 minutes. This data filtering resulted in a total of 1352 sections of 71.2 (range: 10-470) minutes. During the 4 second recording of 10Hz sensor burst, the 1Hz GPS and sensor data recording is not occurring. Thus, to create continuous 1Hz data series, we subsampled the 10Hz data by selecting the first acceleration and magnetometer value of every second during the burst. Linear interpolation was used to fill the 4 s gaps in GPS data occurring during sensor burst. While dead-reckoning provides a more accurate approach to fill data gaps (35), linear interpolation is not expected to introduce a significant bias in the current study due to the overall low proportion of gaps (6±3% percent of all data) and the highly similar distribution at different regions.

### Identifying thermal soaring-gliding

Soaring-gliding events consist of an altitudinal gain phase performed using circular thermal climbing, followed by a gliding phase during which altitude loss occurs (36). Thermal soaring and inter-thermal gliding fundamental movement elements (FMEs) (37) were defined separately in our dataset due to the tendency of the cranes to use a mixture of gliding and flapping flight (1), hence not all thermal soaring phases were followed by clearly definable gliding phases.

To identify soaring phases, we first found all climbing events. Two main types of thermal soaring were observed in our data: (1) classic soaring, or circling flights phases, were identified by a continuous change in heading angle in one direction for at least two full circles during the climb event (97% of all thermal sections) and (2) spring-like soaring pattern, which might be a result of circling with high drift (3% of all thermal sections, 12% of thermal sections over the sea, Fig S2). Those were identified by a continuous change in heading direction using the magnetometer data (Fig. 3). For each local minimum point in the flight height, we found its following local maximum point. If the time between the max and min points was at least 30 seconds, and if for each recorded flight heights 10 seconds apart the later was higher, we considered it a climb. Climbs less than 15 seconds apart were merged. A consequent gliding phase was registered if 80% of the 1 s steps were downwards without circling for at least 30 s and if it followed a soaring phase by less than 60 s. (36).

The mean (±SD) time of thermal soaring was 213 (±116) s, and the mean (±SD) time of gliding was 109 (±82) s. For each segment of thermal soaring, we calculated vertical speed (climb rate), thermal starting and exit heights above terrain, and flapping proportion (see section 2.3.2). When a matching gliding phase was coupled with the soaring phase (61% of the soaring FMEs), time of gliding, ground speed, air speed, vertical speed (sink speed) and flapping rate were calculated (Fig 3).

### Flapping rate

The tri-axial acceleration (in millivolts) was transformed to acceleration (ms^-2^) units using calibration values obtained for each azimuth collected prior to deployment. To obtain vertically aligned acceleration we calculated tri-axial static acceleration and projected the raw acceleration. Flapping frequency was determined by counting the number of detected wingbeats per second, using the high 10 Hz recording during 4 s (cf. 38).

Since commonly used methods (39, 40) for determining the instantaneous flap rate during flight using acceleration or magnitude data are not suitable at the sampling resolution of 1Hz, we developed a model based on a two-layer, fully-connected, feedforward neural network (Python 3.9; TensorFlow 2.8) to estimate flap rate when 10Hz data was not available. The input neurons were the result of applying aggregate functions and calculating the Pearson correlation between axes for acceleration and magnitude data. For model calibration we used 12,580 sections, divided into 80% for model training and 20% for model validation. Models were estimated for multiple configurations and the best model was selected according to the goodness-of-fit of the flap-rate projected from 1Hz data and the one calculated from 10Hz data. The best performing model (R^2^ = 0.91) was based on 5 continuous samples of 1Hz acceleration (magnitude data was dropped in model selection) with the flap rate estimated for the middle sample point. This model was used to calculate flapping rate from 1Hz data for all our tracks (Fig. 2). The mean (±SD) calculated flapping rate was 2.15 (±0.79) flaps s^−1^ during flapping flight and 0.89 (±0.83) flaps s^−1^ during thermal soaring. Flapping rate was converted into flapping ratio to represent flapping proportion during each fundamental movement element (see section 2.3.3).

### Other flight characteristics

To assess thermal soaring under different conditions, we segmented the data into 10-minute sections and only sections lasting 10 full minutes were analyzed. The sections were classified as above sea or above land and only sections for which all points were classified to the same habitat were analyzed. For this analysis only thermal soaring was considered and for each section, the number of thermal soaring FMEs and total time in thermal soaring were recorded.

To understand the differences in crane flight performance at different geographical areas and environmental conditions, we used only coupled soaring-gliding FMEs. We focused on the following decision making proxies: (a) soaring-gliding efficiency calculated as distance traveled when gliding divided by the time spent in the preceding thermal (cf. 41). (b) RAFI to assess risk minimization (36). RAFI index is calculated based on the optimal speed that maximizes the cross-country flight and the best-glide speed that minimizes the sink-rate. These theoretical speeds are calculated based on mean body mass, wing area and wing span for each species using the FLIGHT (version 1.25) software (42). Crane morphological input values (mean body mass = 5.614 kg, wing area = 0.5853 m^2^, wing span = 2.22 m, aspect ratio = 8.42) were obtained from data provided in (43) Lower RAFI index values represent higher airspeed when more risky flight is used, and higher RAFI values represent risk-averse flight behavior with lower sink rates (36). Theoretical RAFI index values, if the bird flies optimally, range between 0 and 1; however, birds are not always using optimal flight and thus the index values are not limited to these values. 3) Flapping ratio during soaring and gliding phases, calculated by dividing the cumulative number of flaps in each FME by the maximal possible number of flaps based on the mean calculated flapping rate during powered flight. This method of flapping ratio calculation probably leads to overestimation of flapping, as a particular second is considered flapping regardless of how many wing flaps were performed in that second (38, 40). However, because the actual number of flaps was unknown and estimated based on a running average over 4 seconds, our calculation is representative of the flapping ratio during the different movement phases.

### Annotating environmental variables

Flight height above ground level was calculated by subtracting from the altitude ASL (44) the ground elevation (ASTER DEM, 1 arc-second spatial resolution) obtained from Env-DATA track annotation service (4). To relate the flight behavior to the time of day, we classified diurnal locations as those collected between sunrise and sunset and regarded the remaining locations as nocturnal.

To identify flight above the Black Sea and Mediterranean Sea we used the Marine Regions shapefile (45) and annotated the corresponding position to the location inside or outside the polygon. Sea crossing was identified if at least one point was located inside the sea polygon. All data tracks were classified to three geographical areas: (a) over Black or Mediterranean Seas, (b) over land north of latitude 32°N and (c) over land south of latitude 32°N. Latitude 32°N was chosen to differentiate tracks above desert or elsewhere because the geographical areas in which the 1Hz data was sampled do not include deserts north of this altitude (SI Appendix, Fig S1).

Atmospheric variables were obtained from the ERA5 hourly data on pressure levels (46) and single levels (47) from 1979 to present, provided by ECMWF. We annotated all crane GPS locations with the following single level variables: mean sea level pressure, air temperature at 2 meters above the surface, sea surface temperature, total cloud cover, and boundary layer height (BLH). Additionally, annotation was done on the 950 millibar pressure level with geopotential height and U- and V-wind components. Since the ERA5 temporal resolution is one hour, the GPS location timestamp was rounded to the nearest hour for the ERA5 annotation. Precipitation data were obtained from the GPCC First Guess Daily Product at 1.0° (48).

For soaring-gliding flight performance analysis, we estimated the wind speed and direction using bird drift in thermals from the horizontal displacement of thermals (49). We chose this estimation because of the coarse resolution of the ERA5 data that may not correctly predict conditions in the small scales of thermal soaring-gliding events. However, this estimate was not possible for 1Hz sections with no thermal soaring; thus, for comparison between soaring and flapping sections, the ERA5 data was used.

### Statistical analysis

To identify factors influencing thermal probability at sea, we modelled the relationship between thermal presence and meteorological predictor variables using binomial Generalized linear-mixed model (GLMM) with the glmer.nd function in the ‘*lme4*’ package (50). Animal ID was included as random factor in all models to account for repeated measures. Before fitting the GLMMs, all continuous predictors were transformed to z-scores to standardize them (51). To rule out collinearity we calculated Pearson’s correlation coefficients (r) between each pair of explanatory variables and selected variables with |r|< 0.7 (52) We compared the full model with a null model (including only random and control variables) using likelihood-ratio tests (ANOVA function set to ‘Chisq’). The response variable was presence/absence of thermal circling, and the predictor meteorological variables included mean tailwinds, mean temperature difference between sea and air (ΔT) and sea level pressure. Mean BLH was correlated with ΔT (r = 0.72, p < 0.001) and excluded from the model. An additional predictor variable was individual age: adults versus birds under the age of three. First year juveniles were not analyzed separately because our high-resolution dataset included only one juvenile.

To compare the proportion of 10 minute sections with and without thermal presence at different geographical regions and between autumn and spring, we applied Chi-squared contingency table analyses to determine the overall effects, followed by post-hoc pairwise comparisons calculated using the Marascuilo procedure (53) which allows comparison of proportion data among several populations. To compare the decision-making proxies between the geographical regions, we applied an aligned rank transformed ANOVA (ART-ANOVA) for non-parametric, factorial analyses with crane ID as a random factor, using the *ARTool* package (54). We conducted within-group comparisons using the *ARTool* pair-wise contrast function and between-group comparisons using Mann-Whitney U-tests with Bonferroni corrected p-values.

We used repeated measures analysis of variance (RM-ANOVA) to compare conditions in the Mediterranean Sea between three one-day timescales prior and one one-day timescale after crane departure for Mediterranean Sea crossing in autumn. A common location (35°17’N, 35°36E) was selected for this analysis because most tracks crossed this area. For each individual, crossing flight mode was set as thermal soaring if at least one 10-minute section with thermal soaring was present during crossing; otherwise, it was set as flapping only. Each meteorological factor (ΔT, sea level pressure, total cloud cover, precipitation, and tailwind) was analyzed separately. Tailwind was calculated in relation to bird flight direction between the point of departure and the point of analysis in sea. We then applied Dunnett’s test (55) to compare the conditions at the departure day with the ones measured at other days (−3, -2, -1 and +1).

## Author contributions

Conceptualization, project administration and supervision: RN

Study design: RN, SP

Fieldwork: SP, YM

Data analysis: SP, DS, NH

Visualization: SP

Funding acquisition: RN, WMG, RCKB, PLK

Writing – original draft: SP, DS, RN

Writing – review & editing: SP, DS, RN, WMG, RCKB, PLK

## Acknowledgments

The authors wish to thank to K. Postelnykh, K. Kondrakova, N.Yesraeli, S. Agmon, Y. Charka, G. Shani, N. Valtzer and F. Argyle for help in fieldwork and trapping. We also thank the members of Nathan’s Movement Ecology lab and especially Y. Bartan and S. Turjeman for their help at various stages of the research. This research was funded by BSF grant 904/2015 to RN and WMG, by GIF grant 999-66.8/2008, ISF grant 2525/16 and JNF/KKL grant 14-093-01-6 to RN, and by NSF grant 1617982 to WMG, RCKB, and PLK. We also acknowledge financial support from the Adelina and Massimo Della Pergola Chair of Life Sciences and the Minerva Center for Movement Ecology to RN.

