## Supplemental Information for "Cranes soar on thermal updrafts behind cold fronts as they migrate across the sea"

### SUPPLEMENTARY FIGURES

**Fig. S1.**

**High resolution data collection regions.** High resolution data recorded in 1 s intervals, once the bird was flying (speed > 2.8 m/s), in pre-set geographical areas along the migration route (red rectangles). Geographical areas used for the analysis are depicted on the figure. 1Hz data shown in yellow lines. The 32N latitude used to differentiate between tracks at the “north” and the “south” is shown in dashed white line.

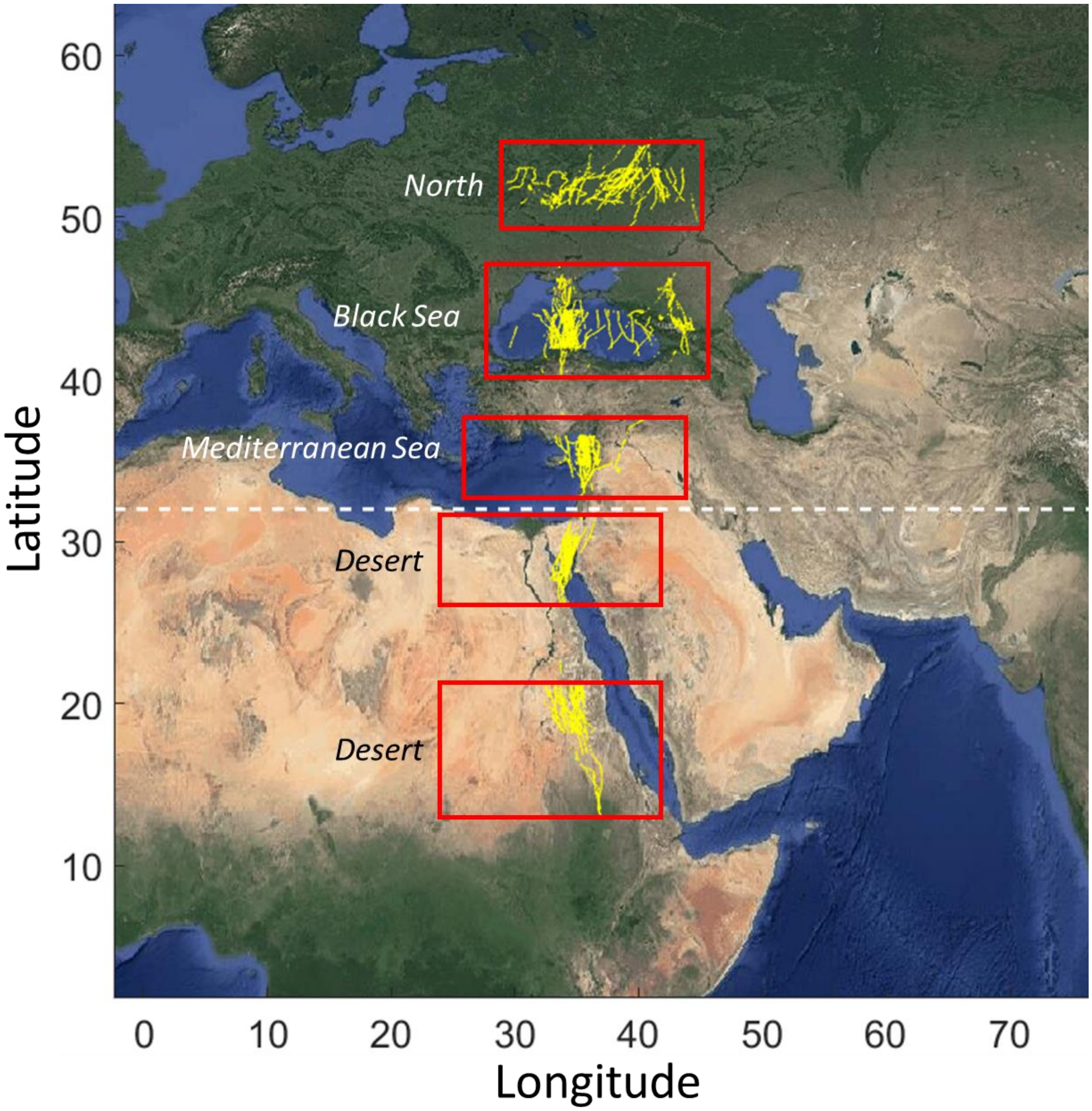

**Fig. S2.**

**Example of thermal soaring of the two main types observed in our data.** (a) classic soaring circling flights phases were identified by a continuous change in heading angle in one direction for at least two full circles during the climb event and (b) spring-like soaring pattern, which might be a result of circling with high drift.

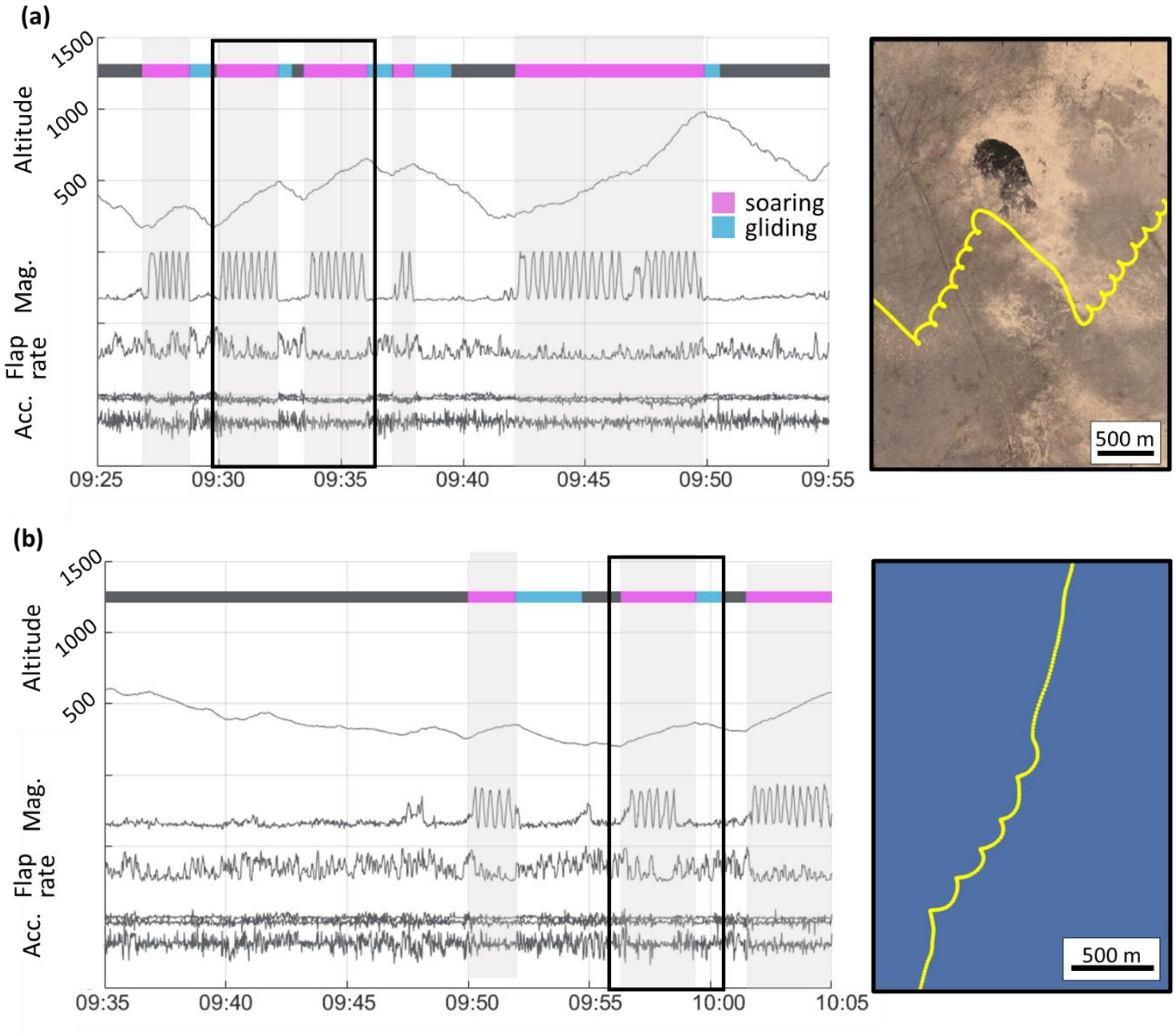

**Fig. S3.**

**Thermal soaring-gliding flight characteristics and performance across three geographical regions.** See Fig. 2 for region classifications and color codes. (a) RAFI calculated based on local wind conditions estimated using the horizontal displacement of thermals. Higher values indicate risk-averse flight behavior; (b) Soaring–gliding efficiency measured by dividing cross-country distance during inter-thermal glides by the time soaring in thermals; (c) height above ground upon exit from thermal climbing. The Boxplots represent averaged individual data of daily means (for days with more than one number of soaring-gliding events) with population median of all samples (sample size: north, N= 182; desert, N= 154; and sea, N= 23) plotted on top. Asterisks indicate significant difference between groups (* $p < 0.05$, ** $p < 0.001$) based on ART-ANOVA with pairwise Bonferroni post-hoc tests.

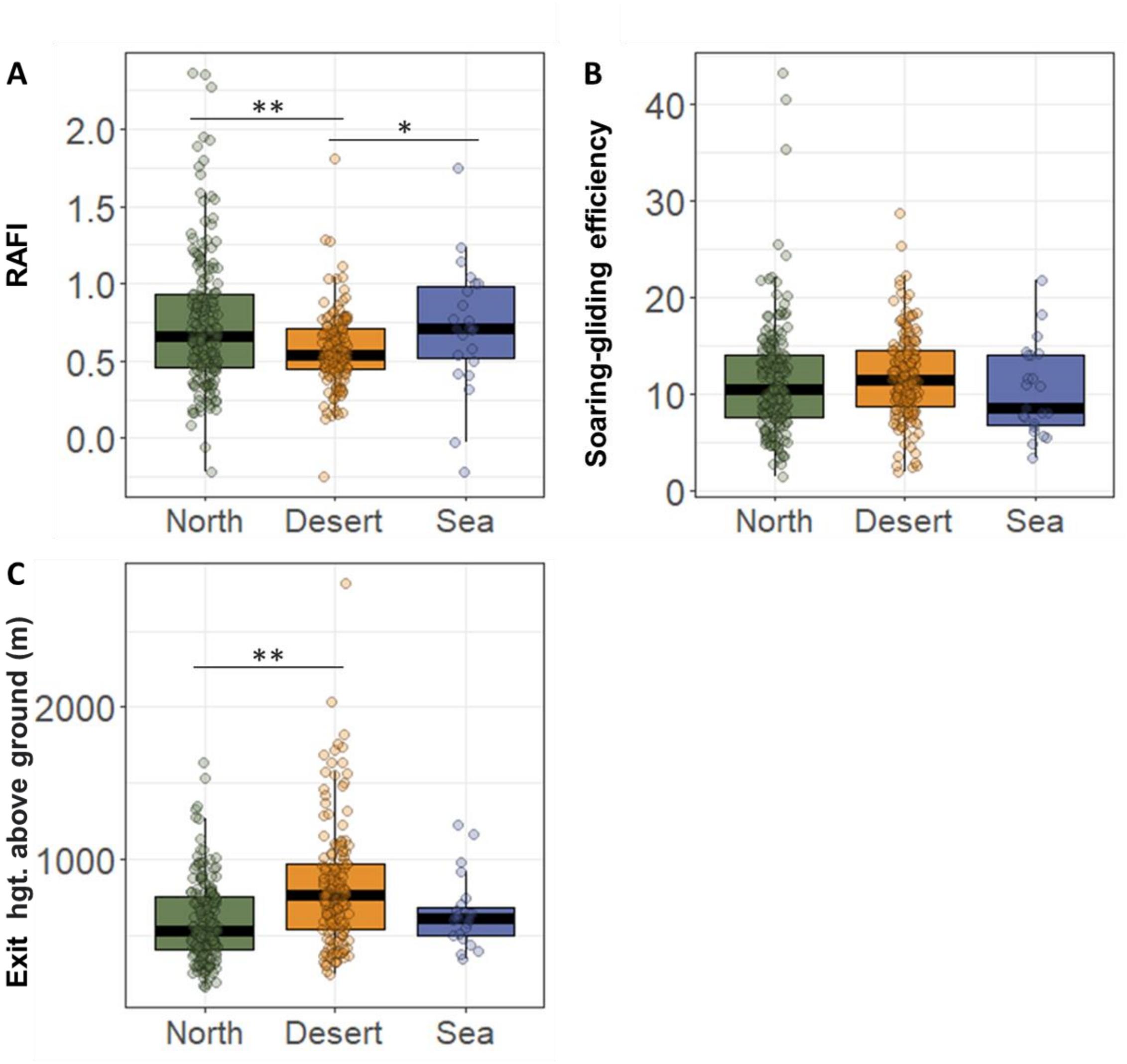

### SUPPLEMENTARY TABLES

**Table S1.**

Species specific morphological attributes used in Fig. 1. Body mass (BM), wing span (WS), wing area (WA), aspect ratio (AR) and wing loading (WL). WL and AR were calculated based on BM, WS and WA provided in the literature. For data from Nudds et al. (2007) WS was missing and thus AR was used.

| Species | BM (kg) | WS (m) | WA($m^2$) | AR | WL ($Nm^{-2}$) | References |
|---|---|---|---|---|---|---|
| *Pelecanus onocrotalus* | 8.504 | 2.91 | 0.955 | 8.863 | 87.32 | (Alerstam et al. 2007) |
| *Ciconia ciconia* | 3.432 | 1.91 | 0.533 | 6.850 | 63.21 | (Alerstam et al. 2007) |
| *Pernis apivorus* | 0.778 | 1.26 | 0.247 | 6.425 | 30.89 | (Alerstam et al. 2007) |
| *Circus pygargus* | 0.291 | 1.09 | 0.135 | 8.801 | 21.15 | (Alerstam et al. 2007) |
| *Pandion haliaetus* | 1.578 | 1.60 | 0.320 | 7.998 | 48.36 | (Alerstam et al. 2007) |
| *Milvus migrans* | 0.815 | 1.52 | 0.281 | 8.237 | 28.50 | (Alerstam et al. 2007) |
| *Circus aeruginosus* | 0.653 | 1.16 | 0.204 | 6.596 | 31.40 | (Alerstam et al. 2007) |
| *Circaetus gallicus* | 1.702 | | 0.412 | 8.551 | 40.52 | (Nudds et al. 2007, Rayner 1988) |
| *Aquila pomari* | 2.015 | 1.80 | 0.513 | 6.316 | 38.53 | (Horvitz et al. 2014) |
| *Aquila nipalensis* | 2.900 | 2.03 | 0.485 | 8.455 | 58.66 | (Horvitz et al. 2014) |
| *Aquila chrysaetos* | 4.198 | | 0.565 | 7.430 | 72.89 | (Nudds et al. 2007, Rayner 1988) |
| *Gyps fulvus* | 7.470 | 2.56 | 0.890 | 7.364 | 82.34 | (Stavros & Nikos 2008) |
| *Coragyps atratus* | 2.12 | 1.44 | 0.349 | 5.942 | 59.59 | (Graves 2017) |
| *Cathartes aura* | 2.14 | 1.74 | 0.453 | 6.652 | 46.39 | (Graves 2017) |
| *Vultur gryphus* | 11.30 | | 0.968 | 7.534 | 114.46 | (Nudds et al. 2007, Rayner 1988) |
| *Gymnogyps californianus* | 10.09 | | 0.975 | 9.484 | 101.55 | (Nudds et al. 2007, Rayner 1988) |
| *Fregata magnificens* | 1.52 | 2.29 | 0.408 | 12.853 | 36.55 | (Pennycuick 1983) |
| *Fregata minor* | 1.23 | 1.98 | 0.354 | 11.075 | 34.09 | (Spear & Ainley 1997) |
| *Diomedea irrorata* | 2.92 | 2.215 | 0.346 | 14.200 | 82.91 | (Spear & Ainley 1997) |
| *Sula nubauxii* | 1.46 | 1.555 | 0.213 | 11.374 | 67.28 | (Spear & Ainley 1997) |
| *Uria anlge* | 1.05 | 0.73 | 0.056 | 9.516 | 183.94 | (Spear & Ainley 1997) |
| *Larus fuscus* | 0.719 | 1.4 | 0.193 | 10.134 | 36.47 | (Alerstam et al. 2007) |
| *Larus argentatus* | 1.14 | 1.34 | 0.197 | 9.124 | 56.93 | (Alerstam et al. 2007) |
| *Merops apiaster* | 0.06 | 0.47 | 0.027 | 8.181 | 21.07 | (Horvitz et al. 2014) |
| *Mareca penelope* | 0.74 | 0.82 | 0.081 | 8.260 | 88.98 | (Alerstam et al. 2007) |
| *As crecca* | 0.35 | 0.59 | 0.043 | 8.133 | 79.76 | (Alerstam et al. 2007) |
| *Corvus corax* | 1.15 | 1.21 | 0.247 | 5.923 | 45.60 | (Alerstam et al. 2007) |
| *Turdus philomelos* | 0.07 | 0.36 | 0.022 | 5.945 | 30.60 | (Alerstam et al. 2007) |
| *Ardea cinerea* | 1.44 | 1.73 | 0.372 | 8.054 | 37.99 | (Alerstam et al. 2007) |
| *Anser anser* | 3.33 | 1.55 | 0.308 | 7.803 | 105.97 | (Alerstam et al. 2007) |

| | | | | | | |
|---|---|---|---|---|---|---|
| *Cygnus cygnus* | 8.69 | 2.29 | 0.605 | 8.675 | 141.01 | (Alerstam et al. 2007) |
| *Limosa lapponica* | 0.32 | 0.73 | 0.052 | 10.248 | 59.99 | (Alerstam et al. 2007) |
| *Grus grus* | 5.61 | 2.22 | 0.586 | 8.417 | 94.06 | (Alerstam et al. 2007) |
| *Falco peregrinus* | 0.79 | 1.02 | 0.126 | 8.277 | 61.58 | (Alerstam et al. 2007) |
| *Falco eleonorae* | 0.39 | 0.95 | 0.104 | 8.670 | 36.47 | (Alerstam et al. 2007) |


**Table S2.**

Calculations of Marascuilo’s procedure for regional differences in proportion of soaring sections (Fig 2B top plot)

| **Difference** | **Value** | **Critical range** | **significance** |
|---|---|---|---|
| North \| Desert | 0.187 | 0.030 | < 0.05 |
| North \| Sea | 0.334 | 0.024 | < 0.05 |
| Desert \| Sea | 0.523 | 0.026 | < 0.05 |

**Table S3.**

Calculations of Marascuilo’s procedure for seasonal differences in proportion of soaring sections (Fig 2B top plot)

| **Difference** | **Value** | **Critical range** | **significance** |
|---|---|---|---|
| North - fall \| North - spring | 0.013 | 0.058 | N.S |
| Desert - fall \| Desert - spring | 0.081 | 0.061 | < 0.05 |
| North - fall \| Desert - fall | 0.135 | 0.058 | < 0.05 |

North - spring | Desert - fall    0.148    0.068    < 0.05

**Table S4.**

Results of the binomial GLMM with bird identity as a random factor. Model estimates for fixed effects on probability of thermal soaring over the sea. SD, standard deviation; z, Z statistics.

| Predictor | Estimate | SD | z | p | mar./con. $R^2$ |
|---|---|---|---|---|---|
| Intercept | -3.84 | 0.41 | -9.29 | <0.001 | 0.31/0.37 |
| Wind speed | 0.06 | 0.03 | 2.26 | 0.024 | |
| Temp. diff. sea-air | 0.59 | 0.05 | 11.90 | <0.001 | |
| sea level pressure | -0.12 | 0.10 | -1.27 | 0.203 | |
| age | -0.44 | 0.39 | -1.14 | 0.255 | |
