## Supplemental Information Data S1 for "Cranes soar on thermal updrafts behind cold fronts as they migrate across the sea"

### Data S1: used for analysis per individual crane, season and year

Number of individuals = 44.

Cumulative time per individual and trip = 677 ± 434 minutes, indicating cumulative length of 1Hz data.

Blank rows left when not applicable due to data absence or gaps in data.

| ID | Year | season | Date of migration start | Date of migration end | Age | sex | Mig. duration (days) | N. mig. bouts | 1Hz time (minutes) |
|---|---|---|---|---|---|---|---|---|---|
| 16113 | 2018 | fall | 9-Oct-18 | 7-Nov-18 | Adult | Male | 30 | 4 | 87.9 |
| 161152 | 2018 | fall | 17-Sep-18 | 27-Oct-18 | Subadult | Male | 41 | 6 | 330.2 |
| 161152 | 2019 | fall | 14-Oct-19 | 20-Oct-19 | Adult | Male | 7 | 7 | 21.2 |
| 161152 | 2020 | spring | 1-Mar-20 | 22-Aug-20 | Adult | Male | | 4 | 303.4 |
| 161161 | 2018 | fall | 9-Oct-18 | 6-Nov-18 | Adult | Female | 29 | 2 | 85.6 |
| 170511 | 2019 | spring | 1-Mar-19 | 22-Mar-19 | Adult | Male | 22 | 5 | 853.5 |
| 170521 | 2018 | fall | 9-Oct-18 | 7-Nov-18 | Adult | Male | 30 | 12 | 1173.8 |
| 170521 | 2019 | fall | 5-Oct-19 | 10-Nov-19 | Adult | Male | 37 | 15 | 929.4 |
| 170521 | 2019 | spring | 26-Feb-19 | 18-Apr-19 | Adult | Male | 52 | 18 | 1165.8 |
| 170521 | 2020 | fall | 17-Oct-20 | 4-Nov-20 | Adult | Male | 19 | 13 | 833 |
| 170521 | 2020 | spring | 23-Feb-20 | 8-Apr-20 | Adult | Male | 46 | 25 | 1432.4 |
| 170521 | 2021 | spring | 25-Feb-21 | 1-Apr-21 | Adult | Male | | 16 | 1237.3 |
| 170531 | 2018 | fall | 5-Oct-18 | 29-Oct-18 | Adult | Female | 25 | 7 | 359.1 |
| 170541 | 2018 | fall | 5-Sep-18 | 24-Oct-18 | Adult | Male | 50 | 8 | 552.7 |
| 170541 | 2019 | fall | 23-Sep-19 | 20-Oct-19 | Adult | Male | 28 | 7 | 411.2 |
| 170541 | 2020 | fall | 20-Sep-20 | 6-Nov-20 | Adult | Male | 48 | 13 | 995.3 |
| 170541 | 2021 | spring | 23-Feb-21 | 17-Apr-21 | Adult | Male | | 20 | 1132.8 |
| 170551 | 2018 | fall | 10-Oct-18 | 2-Nov-18 | Adult | Female | 24 | 5 | 1015.7 |
| 170581 | 2018 | fall | 11-Oct-18 | 1-Nov-18 | Adult | Male | 22 | 13 | 921.6 |
| 170581 | 2019 | fall | 14-Oct-19 | 9-Nov-19 | Adult | Male | 27 | 12 | 657.6 |
| 170581 | 2019 | spring | 13-Mar-19 | 9-Apr-19 | Adult | Male | 28 | 15 | 913.1 |
| 170581 | 2020 | fall | 17-Oct-20 | 18-Nov-20 | Adult | Male | 33 | 12 | 1100.8 |
| 170581 | 2020 | spring | 8-Mar-20 | 11-Apr-20 | Adult | Male | 35 | 17 | 1002.7 |
| 170581 | 2021 | spring | 7-Mar-21 | 6-Apr-21 | Adult | Male | | 18 | 1310.4 |
| 170591 | 2018 | fall | 9-Oct-18 | 27-Oct-18 | Adult | Male | | 8 | 579.5 |
| 170591 | 2019 | fall | 13-Oct-19 | 18-Oct-19 | Adult | Male | 6 | 5 | 427.4 |
| 170591 | 2019 | spring | 6-Mar-19 | 22-Mar-19 | Adult | Male | 17 | 7 | 53.1 |
| 170761 | 2018 | fall | 9-Oct-18 | 31-Oct-18 | Adult | Male | 23 | 11 | 806.9 |
| 170761 | 2019 | fall | 14-Oct-19 | 20-Nov-19 | Adult | Male | 38 | 12 | 1113.5 |
| 170761 | 2019 | spring | 23-Feb-19 | 1-Apr-19 | Adult | Male | 38 | 20 | 2016.5 |
| 170761 | 2020 | spring | 28-Feb-20 | 31-Mar-20 | Adult | Male | 33 | 20 | 1031.5 |
| 170771 | 2018 | fall | 5-Oct-18 | 24-Oct-18 | Subadult | Male | 20 | 5 | 607.5 |
| 170791 | 2019 | fall | 14-Oct-19 | 30-Oct-19 | Adult | Male | 17 | 12 | 360.5 |
| 170791 | 2020 | fall | 17-Oct-20 | 8-Nov-20 | Adult | Male | 23 | 14 | 520 |
| 170791 | 2020 | spring | 24-Feb-20 | 30-Mar-20 | Adult | Male | 36 | 18 | 2314.4 |
| 170791 | 2021 | spring | 24-Feb-21 | 10-Apr-21 | Adult | Male | | 20 | 1589.1 |
| 170801 | 2018 | fall | 9-Oct-18 | 16-Oct-18 | Adult | Male | 8 | 5 | 664.8 |
| 170801 | 2019 | spring | 8-Mar-19 | 23-Mar-19 | Adult | Male | 16 | 11 | 495.4 |
| 170801 | 2020 | fall | 17-Oct-20 | 7-Nov-20 | Adult | Male | 22 | 6 | 596.3 |
| 170801 | 2021 | spring | 16-Mar-21 | 29-Mar-21 | Adult | Male | | 7 | 235.3 |
| 170811 | 2019 | fall | 1-Sep-19 | 20-Nov-19 | Adult | Male | | 5 | 332 |
| 170811 | 2020 | fall | 23-Sep-20 | 19-Nov-20 | Adult | Male | | 13 | 766.7 |
| 170811 | 2020 | spring | 8-Mar-20 | 19-Apr-20 | Adult | Male | 43 | 13 | 420 |

| | | | | | | | | | |
|---|---|---|---|---|---|---|---|---|---|
| 170811 | 2021 | spring | 1-Mar-21 | 9-Apr-21 | Adult | Male | | 19 | 1102.9 |
| 170821 | 2018 | fall | 9-Oct-18 | 2-Nov-18 | Subadult | Male | 25 | 13 | 729.4 |
| 170821 | 2019 | fall | 5-Oct-19 | 22-Nov-19 | Adult | Male | 49 | 12 | 726.6 |
| 170821 | 2020 | fall | 17-Oct-20 | 21-Oct-20 | Adult | Male | 5 | 6 | 183.5 |
| 170821 | 2020 | spring | 5-Mar-20 | 13-Apr-20 | Adult | Male | 40 | 18 | 600 |
| 170831 | 2018 | fall | 9-Oct-18 | 14-Oct-18 | Adult | Male | 6 | 6 | 208.3 |
| 170831 | 2019 | fall | 14-Oct-19 | 19-Oct-19 | Adult | Male | 6 | 5 | 220.6 |
| 170861 | 2018 | fall | 5-Oct-18 | 27-Oct-18 | Adult | Male | 23 | 5 | 862.6 |
| 170861 | 2019 | fall | 5-Oct-19 | 3-Nov-19 | Adult | Male | 30 | 5 | 553.7 |
| 170861 | 2019 | spring | 6-Mar-19 | 25-Mar-19 | Adult | Male | 20 | 7 | 868.6 |
| 170861 | 2020 | fall | 17-Oct-20 | 8-Nov-20 | Adult | Male | | 3 | 244 |
| 170861 | 2020 | spring | 3-Mar-20 | 29-Mar-20 | Adult | Male | | 6 | 593.8 |
| 170861 | 2021 | spring | 26-Feb-21 | 1-Apr-21 | Adult | Male | | 6 | 412.7 |
| 170871 | 2018 | fall | 5-Oct-18 | 7-Nov-18 | Subadult | Male | 34 | 7 | 1434.4 |
| 170881 | 2019 | fall | 14-Oct-19 | 19-Oct-19 | Adult | Female | 6 | 5 | 474.1 |
| 170891 | 2018 | fall | 9-Oct-18 | 3-Nov-18 | Adult | Male | 26 | 12 | 1367.1 |
| 170891 | 2019 | fall | 5-Oct-19 | 1-Nov-19 | Adult | Male | 28 | 13 | 1176.7 |
| 170891 | 2019 | spring | 12-Mar-19 | 11-May-19 | Adult | Male | 61 | 21 | 655.2 |
| 170891 | 2020 | fall | 17-Oct-20 | 22-Oct-20 | Adult | Male | 6 | 6 | 260.3 |
| 170891 | 2020 | spring | 5-Mar-20 | 9-Apr-20 | Adult | Male | 36 | 17 | 1173.9 |
| 170891 | 2021 | spring | 29-Jan-21 | 31-Mar-21 | Adult | Male | | 7 | 386.1 |
| 170911 | 2018 | fall | 9-Oct-18 | 1-Nov-18 | Adult | Male | 24 | 11 | 1101.2 |
| 170911 | 2019 | fall | 14-Oct-19 | 4-Nov-19 | Adult | Male | 22 | 12 | 995.4 |
| 170911 | 2019 | spring | 25-Feb-19 | 21-Apr-19 | Adult | Male | 56 | 19 | 597.6 |
| 170911 | 2020 | fall | 17-Oct-20 | 3-Nov-20 | Adult | Male | 18 | 14 | 1045.6 |
| 170911 | 2020 | spring | 29-Feb-20 | 3-Apr-20 | Adult | Male | 35 | 14 | 1377.2 |
| 170911 | 2021 | spring | 25-Feb-21 | 1-Apr-21 | Adult | Male | | 16 | 1693 |
| 170931 | 2018 | fall | 9-Oct-18 | 13-Oct-18 | Adult | Female | 5 | 3 | 107.8 |
| 170931 | 2019 | fall | 14-Oct-19 | 19-Oct-19 | Adult | Female | 6 | 6 | 445 |
| 170931 | 2020 | fall | 17-Oct-20 | 15-Dec-20 | Adult | Female | 60 | 13 | 417.5 |
| 170931 | 2020 | spring | 9-Mar-20 | 28-Mar-20 | Adult | Female | 20 | 9 | 1870.7 |
| 170931 | 2021 | spring | 3-Mar-21 | 10-Apr-21 | Adult | Female | | 18 | 1326 |
| 170951 | 2018 | fall | 9-Oct-18 | 14-Oct-18 | Adult | Male | 6 | 5 | 349.1 |
| 170951 | 2019 | fall | 14-Oct-19 | 24-Oct-19 | Adult | Male | 11 | 7 | 551.1 |
| 170961 | 2018 | fall | 9-Oct-18 | 16-Oct-18 | Adult | Male | 8 | 11 | 412 |
| 170961 | 2019 | fall | 5-Oct-19 | 4-Nov-19 | Adult | Male | 31 | 8 | 995.6 |
| 170991 | 2018 | fall | 5-Oct-18 | 11-Oct-18 | Adult | Male | 7 | 6 | 292.9 |
| 170991 | 2019 | fall | 5-Oct-19 | 25-Oct-19 | Adult | Male | | 5 | 641 |
| 170991 | 2020 | fall | 17-Oct-20 | 24-Oct-20 | Adult | Male | 8 | 7 | 502.1 |
| 170991 | 2020 | spring | 26-Mar-20 | 3-Apr-20 | Adult | Male | 9 | 6 | 35.3 |
| 170991 | 2021 | spring | 11-Mar-21 | 28-Mar-21 | Adult | Male | 18 | 8 | 767.8 |
| 171001 | 2018 | fall | 9-Oct-18 | 4-Nov-18 | Adult | Female | 27 | 13 | 858.5 |
| 171021 | 2018 | fall | 30-Sep-18 | 15-Oct-18 | Adult | Female | 16 | 7 | 797.6 |
| 171021 | 2019 | fall | 28-Sep-19 | 19-Oct-19 | Adult | Female | 22 | 6 | 239 |
| 171021 | 2020 | fall | 8-Oct-20 | 28-Oct-20 | Adult | Female | 21 | 13 | 796.7 |
| 171021 | 2021 | spring | 26-Jan-21 | 5-Apr-21 | Adult | Female | | 10 | 231.9 |
| 171031 | 2018 | fall | 17-Sep-18 | 17-Nov-18 | Adult | Male | 62 | 14 | 1589.1 |
| 171031 | 2019 | spring | 8-Mar-19 | 8-Apr-19 | Adult | Male | 32 | 21 | 289.4 |
| 171041 | 2018 | fall | 5-Oct-18 | 31-Oct-18 | Adult | Male | 27 | 13 | 430.4 |
| 171041 | 2019 | fall | 5-Oct-19 | 12-Nov-19 | Adult | Male | | 13 | 1105.3 |

| | | | | | | | | | |
|---|---|---|---|---|---|---|---|---|---|
| 171041 | 2019 | spring | 27-Feb-19 | 25-Mar-19 | Adult | Male | 27 | 16 | 815.1 |
| 171041 | 2020 | spring | 23-Feb-20 | 19-Mar-20 | Adult | Male | 26 | 16 | 750 |
| 171042 | 2020 | fall | 17-Oct-20 | 23-Oct-20 | Adult | Unknown | 7 | 9 | 32.5 |
| 171051 | 2018 | fall | 9-Oct-18 | 20-Oct-18 | Juvenile | Female | | 6 | 792.4 |
| 1819491 | 2018 | fall | 9-Oct-18 | 23-Oct-18 | Adult | Male | 15 | 7 | 279.7 |
| 1819491 | 2019 | fall | 14-Oct-19 | 19-Oct-19 | Adult | Male | 6 | 7 | 281.7 |
| 1819491 | 2019 | spring | 8-Mar-19 | 1-Apr-19 | Adult | Male | 25 | 5 | 724.4 |
| 1819491 | 2020 | spring | 26-Feb-20 | 30-Mar-20 | Adult | Male | 34 | 9 | 855.1 |
| 1819501 | 2019 | fall | 14-Oct-19 | 18-Nov-19 | Adult | Male | 36 | 10 | 663.6 |
| 1819501 | 2020 | fall | 17-Oct-20 | 23-Oct-20 | Adult | Male | 7 | 7 | 413.5 |
| 1819501 | 2020 | spring | 16-Mar-20 | 3-Apr-20 | Adult | Male | 19 | 9 | 386.2 |
| 1819501 | 2021 | spring | 15-Mar-21 | 29-Mar-21 | Adult | Male | | 7 | 504.6 |
| 1819511 | 2018 | fall | 9-Oct-18 | 27-Oct-18 | Adult | Female | 19 | 4 | 977.5 |
| 1819511 | 2019 | fall | 14-Oct-19 | 18-Nov-19 | Adult | Female | 36 | 10 | 604.8 |
| 1819521 | 2019 | fall | 5-Oct-19 | 18-Oct-19 | Subadult | Female | 14 | 7 | 640.1 |
| 1819531 | 2018 | fall | 9-Oct-18 | 27-Oct-18 | Juvenile | Male | | | 382.9 |
| 1819531 | 2019 | fall | 14-Oct-19 | 26-Oct-19 | Subadult | Male | 13 | 7 | 223.4 |
| 1819531 | 2020 | fall | 17-Oct-20 | 8-Nov-20 | Adult | Male | 23 | 6 | 447.6 |
| 1819531 | 2021 | spring | 14-Mar-21 | 3-Apr-21 | Adult | Male | | 7 | 251 |
| 1819541 | 2018 | fall | 9-Oct-18 | 28-Oct-18 | Adult | Female | 20 | 7 | 608.1 |
| 1819541 | 2019 | fall | 14-Oct-19 | 30-Oct-19 | Adult | Female | 17 | 8 | 700 |
| 1819541 | 2019 | spring | 6-Mar-19 | 31-Mar-19 | Adult | Female | 26 | 10 | 538.2 |
| 1819541 | 2020 | fall | 17-Oct-20 | 23-Oct-20 | Adult | Female | 7 | 8 | 312 |
| 1819541 | 2020 | spring | 17-Mar-20 | 7-Apr-20 | Adult | Female | 22 | 8 | 461.5 |
| 1819551 | 2019 | fall | 14-Oct-19 | 30-Oct-19 | Adult | Male | 17 | 8 | 772.5 |
| 1819551 | 2020 | fall | 17-Oct-20 | 23-Oct-20 | Adult | Male | 7 | 8 | 298.2 |
| 1819551 | 2020 | spring | 16-Mar-20 | 3-Apr-20 | Adult | Male | 19 | 7 | 439.2 |
| 1819551 | 2021 | spring | 7-Mar-21 | 6-Apr-21 | Adult | Male | | 5 | 228.8 |
| 1819561 | 2018 | fall | 30-Sep-18 | 12-Oct-18 | Adult | Male | 13 | 7 | 66.3 |
| 1819561 | 2019 | fall | 5-Oct-19 | 19-Oct-19 | Adult | Male | 15 | 7 | 683.7 |
| 1819561 | 2020 | fall | 17-Oct-20 | 24-Oct-20 | Adult | Male | 8 | 7 | 328.7 |
| 1819561 | 2020 | spring | 27-Feb-20 | 29-Mar-20 | Adult | Male | 32 | 10 | 999.4 |
| 1819561 | 2021 | spring | 6-Mar-21 | 1-Apr-21 | Adult | Male | | 6 | 279.1 |
| 1819572 | 2020 | fall | 4-Oct-20 | 22-Oct-20 | Adult | Unknown | 19 | 9 | 535.7 |
| 1819581 | 2020 | fall | 4-Oct-20 | 4-Nov-20 | Adult | Unknown | 32 | 15 | 1380.1 |
| 1819581 | 2021 | spring | 2-Feb-21 | 17-Apr-21 | Adult | Unknown | | 23 | 1063 |
| 1819591 | 2020 | fall | 17-Oct-20 | 24-Oct-20 | Adult | Unknown | 8 | 8 | 331.2 |
| 1819591 | 2021 | spring | 17-Mar-21 | 6-Apr-21 | Adult | Unknown | | 7 | 397 |
| 1819611 | 2019 | fall | 3-Sep-19 | 31-Oct-19 | Adult | Male | 59 | 6 | 241.4 |
| 1819611 | 2020 | fall | 4-Oct-20 | 9-Nov-20 | Adult | Male | 37 | 6 | 454.8 |
| 1819611 | 2020 | spring | 23-Mar-20 | 14-Apr-20 | Adult | Male | 23 | 10 | 77.2 |
| 1819611 | 2021 | spring | 7-Mar-21 | 5-May-21 | Adult | Male | | 13 | 345.1 |
